# Species synonyms depict changing but taxon-independent taxonomic praxis

**DOI:** 10.1101/2025.04.28.651127

**Authors:** Vikram Iyer, Avanthika Prasad

**Affiliations:** Department of Biology, Trivedi School of Biosciences, Ashoka University, Ashoka University, Sonipat, Haryana, India; Department of Biology, Indian Institute of Science Education and Research (IISER), Pune, Pune, Maharashtra, India

**Author notes:** Corresponding author: Vikram Iyer.

## Abstract

What constitutes a single species is ultimately arbitrary, resulting in idiosyncrasies in taxonomic praxis determining species counts. One consequence of this arbitrariness is taxonomic synonymization, wherein variation determined not unique enough to constitute distinct species gets merged into one. Synonymization is non-random and subject to taxon and trait-specific biases. We explore these trends using a comprehensive dataset of the world’s marine species for all large phyla and Animalia. We match all accepted species-names from the present day with their synonyms and calculate metrics of synonymization including the time lag between the years of descriptions of a species and its newest synonym. We find that although species description trends across phyla show no consistent, discernible trends, the fraction of names that gets synonymised remains initially constant across phyla, with a subsequent gradual decrease. Most accepted species described around the early 20th century possess synonyms created in the same year, indicating synonyms arising due to new name combinations, suggesting a distinct period of taxonomic praxis. This period was followed by a period of most species descriptions lacking synonyms on moving towards the present. Across taxa, we find the mean lag to be strongly negatively correlated to time, with time not being a limiting factor determining this trend. We also find the number of synonyms per accepted name to also be strongly negatively correlated to time in a taxon-independent manner. Our study considers when the diversity characterising species in the form of once-unique species-names becomes known to science. Our results suggest that across taxa, species described more recently encompass less biological variability, indicating a gradual change in how morphologically delineated species which comprise the vast majority of species described across time are described.

## Introduction

The delineation of species boundaries, that is, the range of variation that characterises a single species, is ultimately arbitrary (De Queiroz, 2007). A consequence of this is that idiosyncrasies in taxonomic praxis can determine species counts. For example, using phylogenetic species concepts results in higher species counts relative to morphological species delimitation (Agapow et al., 2004), and persistent taxon-specific practices of lumping together morphological variation into infraspecific taxa, or splitting them into distinct species determine species counts (Isaac et al., 2004). Taxonomic synonymization is another consequence of the arbitrariness of species delimitation. Describing a species generates the testable hypothesis that the holotype individuals formally attached to the species name represent a distinct group, which, if falsified, results in their synonymization with a pre-existing species name (Gaston and Mound, 1997). These pending synonymizations can result in inflated species-counts until rectified by taxonomic revision (Alroy, 2002).

When two formerly unique species are synonymised into one, one of the species-names becomes a synonym of the other following the Principle of Priority (Article 23, “The Code Online | International Commission on Zoological Nomenclature”). Synonymization occurs non-randomly, with species attributes like body size and range size correlating with synonym number (Guedes et al., 2025; Jones et al., 2012). However, synonym count also correlates with indices of taxonomic praxis like how intensively a taxon is studied (Appeltans et al., 2012). These suggest that synonymization displays taxon-specific trends, driven by taxon-specific differences in taxonomic praxis and traits like motility. We explore these trends using the World Register of Marine Species dataset—a comprehensive dataset of the world’s marine species-names (Appeltans et al., 2012; WoRMS Editorial Board, 2024). We choose to use a marine dataset because most large and biologically diverse animal phyla are mostly or completely arine, and the taxonomists describing them likely represent a large and diverse group as well.

## Methods

We used the World Register of Marine Species (WoRMS, https://www.marinespecies.org/; WoRMS Editorial Board, 2024; accessed 25-07-2023, doi:10.14284/170) which is a comprehensive and regularly updated species-name database of all marine organisms. From this, we identified large phyla containing more than 10,000 described species-names. This includes both currently accepted species names as well as synonyms. Next, we isolated the year each species-name was described in from the complete binomial name which includes describing authors’ names. Describing authors are listed in the field “scientificNameAuthorship” in the WoRMS dataset. If two sets of authors and years were listed, the earlier description was considered. These constitute only 1.90% of the complete Animalia dataset of accepted and unaccepted species names.

For each accepted species, we queried all synonyms listing it as its accepted name and matched each species to its synonyms. For each accepted name described in a given year, we also calculated the time lag in years between the year it was described and the year its newest synonym was described. The first described name of a species remains its accepted name and takes precedence over synonym-names as per the Principle of Priority (Article 23; “The Code Online | International Commission on Zoological Nomenclature”) which applies in the overwhelming majority of cases. As an example, *Chama broderipi* Reeve, 1846 and *Chama segmentina* Clessin, 1889 are today synonyms of *Chama pacifica* Broderip, 1835, with *C. pacifica* being the accepted species name since it was described first. This lag acts as a conservative estimate for the time taken for all the biological diversity—in terms of once-unique species—characterising a species in the present day (2023 CE) to become known to science. For accepted names whose newest synonym bore the same year of publication, we also identified synonyms with similar specific epithets. A specific epithet was regarded as “similar” if its first four letters were the same for the accepted name and its newest synonym. A major source of such similar specific epithets is when taxonomic revisions occur at the genus level and new combinations are created (Article 34.2; Article 48, “The Code Online | International Commission on Zoological Nomenclature”).

For each year starting from 1757 CE, we calculated the proportion of accepted names to the total names described in that year, and the mean number of synonyms per each accepted name per year. For these, we also calculated Spearman correlation coefficients for each 30-year period starting from 1757 CE. This acts as an index of the monotonicity of the relationship observed. All data were analysed and plotted in Python ver. 3.12.5. Spearman correlations were calculated using the SciPy function “scipy.stats.spearmanr” (Virtanen et al., 2020) and straight lines and confidence intervals were fit and plotted to the data using the Seaborn function “seaborn.regplot” (Waskom, 2021) after square-root transformation.

## Results

### Species-description trends vary widely across taxa

Moving averages of accepted species-names revealed that there is no consistent trend across phyla (Figure 1), with larger sliding window lengths being less sensitive to outliers. Linnaeus’ continuing legacy could be seen in high starting values at 1757 CE for phyla which are charismatic such as Chordata and for phyla with persistent skeletons such as Mollusca. On the other hand, for soft-bodied phyla such as Platyhelminthes, Nematoda, and Annelida, species descriptions before around 1850 CE were very low. The largely microscopic Foraminifera also presented low description rates in this period.

**Figure 1:**
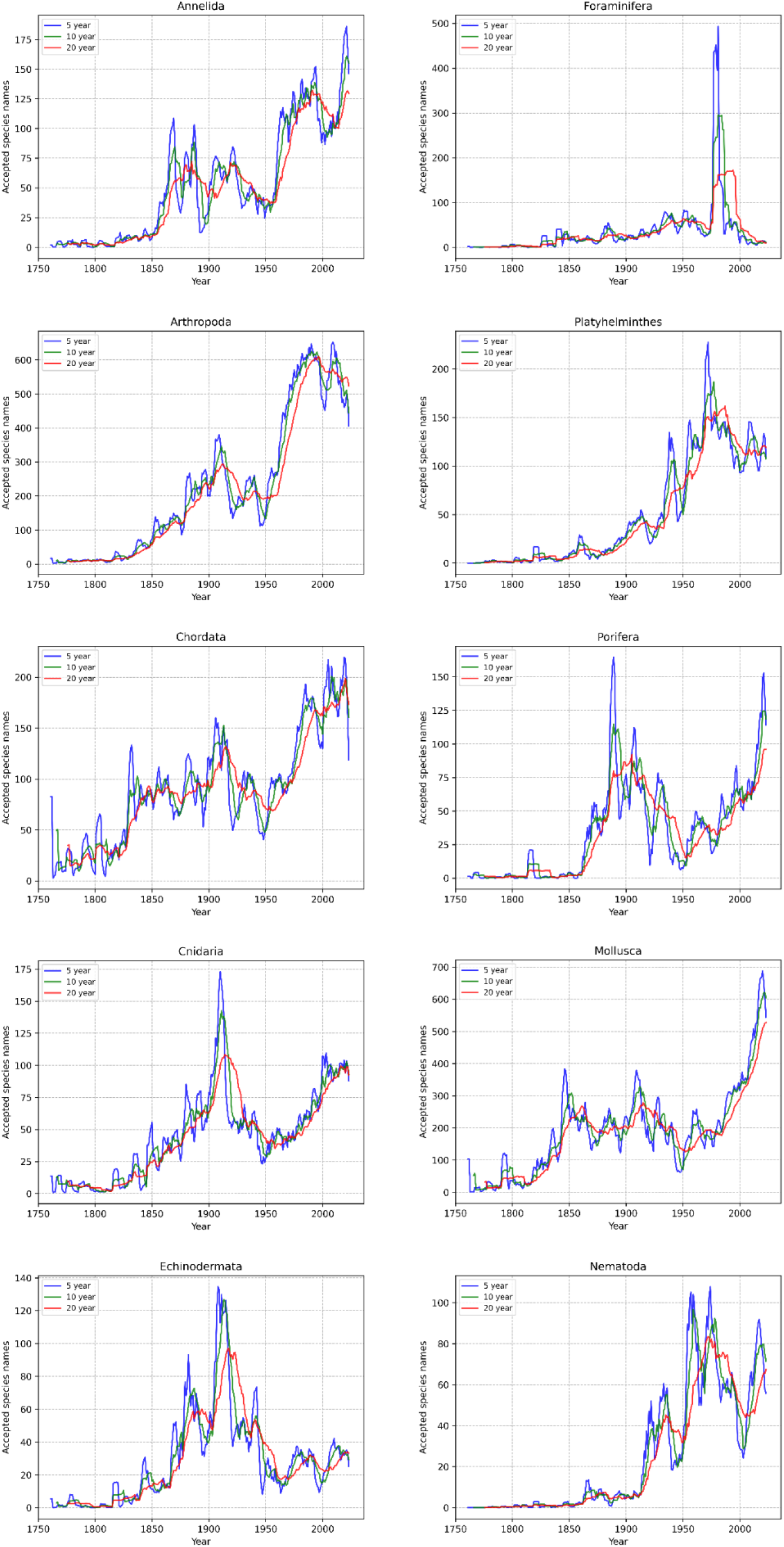
5, 10, and 20 year moving averages of the number of accepted species names with their year of publication for each phylum. For each year, the graph indicates how many species names which are accepted in 2023 were published in that year.

Large and prominent phyla like Arthropoda, Mollusca, Chordata, and Cnidaria displayed a continuously increasing trend in species described from the 1950s onward, with Echinodermata as a notable exception. Outlier years where a large number of species are described in a single year became common from around 1850 to 1950 but were rare thereafter except in chordates.

### Synonymisation trends reveal changing taxonomic praxis consistently across taxa

For kingdom Animalia, the proportion of all names published in a given year which are valid today increased with time (Figure 2a). This trend held for all large phyla as well (Supplementary figure S1). This is reflected in increasing Spearman correlation coefficients for non-overlapping time bins starting from 1757 in a taxon-independent manner (Figure 2b). Accepted species-names with at least one synonym decreased in proportion from the early 20th century onwards, and the vast increase in named species in the second half of the 20th century was driven by species lacking any synonym (Figure 2c). Accepted species-names for which their newest synonym was published after a time-lag decreased in abundance from the late 19th century onwards. Amongst accepted names with no time-lag in publication, the subset of accepted names for which the newest synonym had a specific epithet similar to theirs displayed similar trends to the overall pattern of zero-lag synonyms with time (Figure 2d).

**Figure 2:**
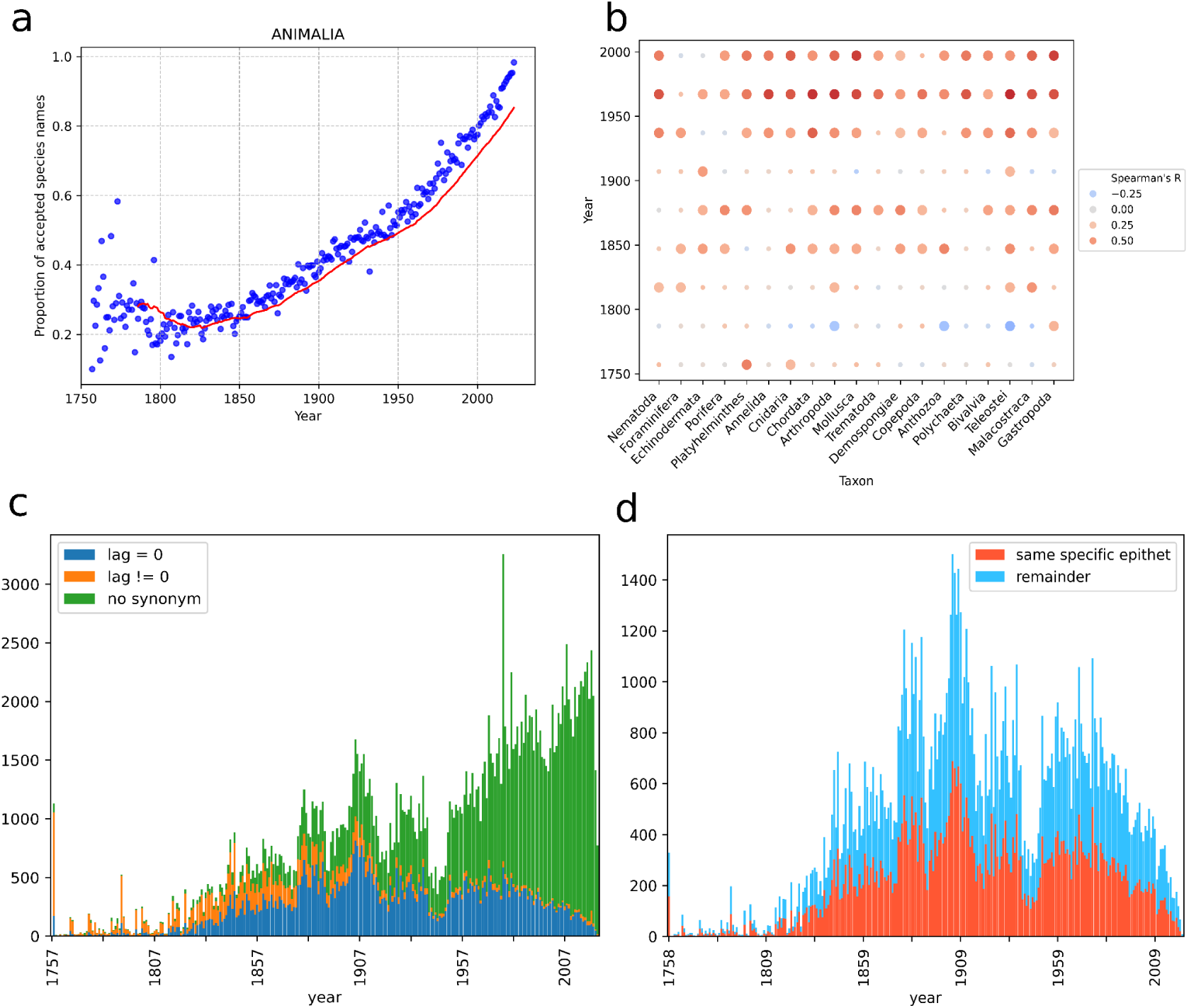
Synonymization with time for Animalia. **A:** Scatterplot depicting the fraction of all species-names published per year which are accepted today. Points represent values for a year and the red line displays the 20 year moving average. **B:** Spearman correlation coefficients between year and the proportion of accepted species names for all large taxa for each 50 year period starting from 1757. Large circles represent significant (p < 0.1) correlations. **C–D:** Stacked histograms depicting the number of accepted species-names (c) in present day grouped by the lag between the accepted species-name and its newest synonym. “Remainder” depicts accepted species-names without a synonym, (d) with a lag time of zero between the name and its newest synonym, grouped by whether or not the specific epithets are similar between them. “Remainder” depicts accepted species-names with their newest synonym bearing a dissimilar specific epithet.

The mean lag times between the accepted names published in a year and their newest synonyms were strongly negatively correlated to time after square-root transformation (Spearman R = – 0.88, p < 0.0001) and displayed a linear trend (Figure 3a). These mean lag values were smaller than the time remaining till present day until after approximately 2000 CE. Overall, the same trends held across large phyla (Supplementary figure S2). The mean number of synonyms per accepted name was also strongly negatively correlated to time after square-root transformation (Spearman R = – 0.94, p < 0.0001) and displayed a linear trend as well (Figure 3b) across all large phyla (Supplementary figure S3).

**Figure 3:**
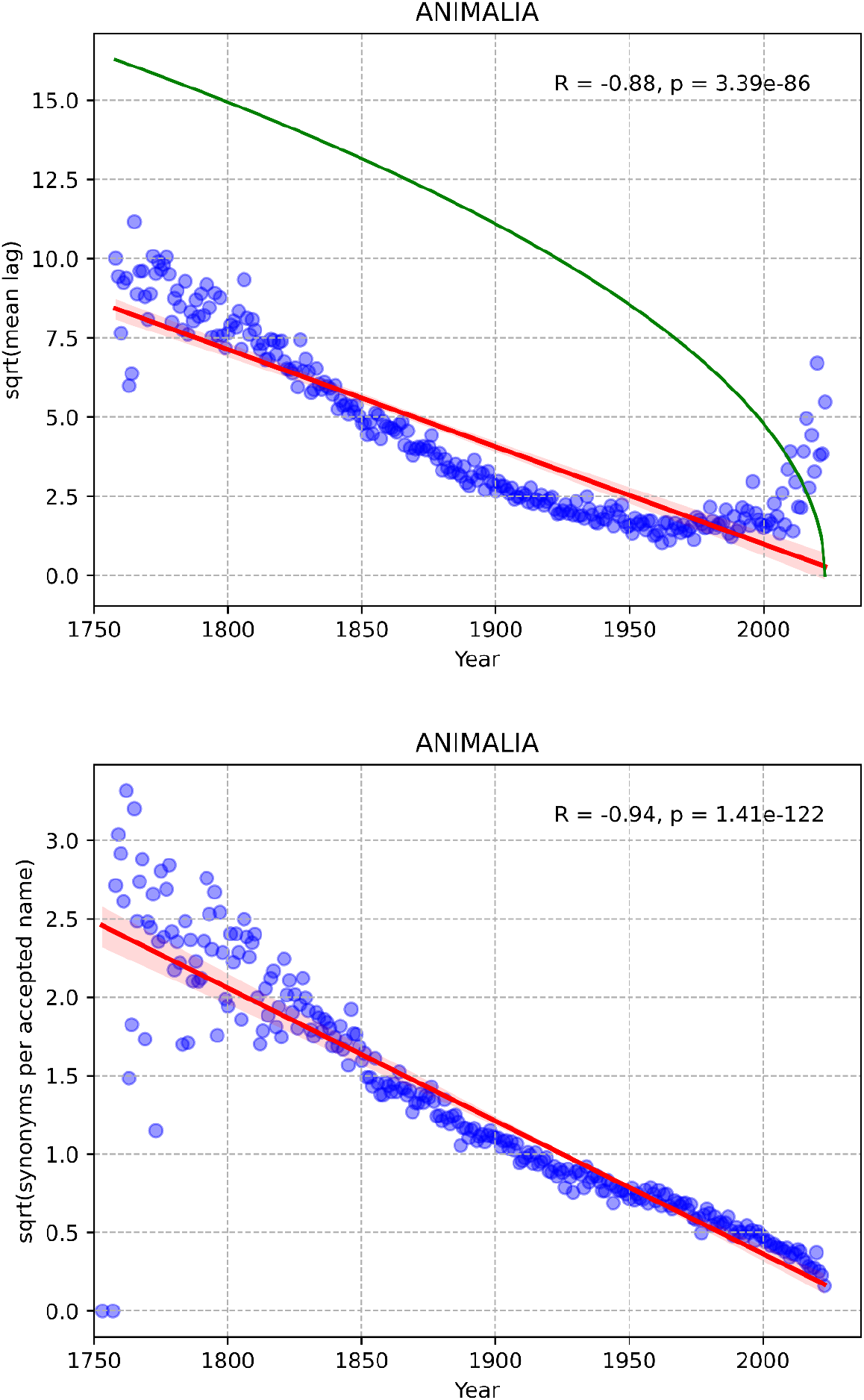
Metrics of synonymization with time for Animalia. Above: Scatterplot depicting the mean lag value between the years of description of the newest synonym of the accepted species names published in a year. Points represent lag values. The red line represents a straight line fit to the data and red bands indicate 95% confidence intervals. R = Spearman correlation coefficient. Lag values are square-root transformed before plotting. The green line represents the number of years till present day. Below: Scatterplot depicting the mean number of synonyms for each accepted species-name published with time. Values are square-root transformed before plotting. Points represent the mean number of synonyms. The red line represents a straight line fit to the data and red bands indicate 95% confidence intervals. R = Spearman correlation coefficient.

## Discussion

A once-distinct species name which later gets synonymised encompasses biological variability that is not distinct enough to warrant its classification as a unique species. We do not consider when two species-names become synonymised. Instead, we consider when all the diversity characterising a species—represented as different once-unique species which eventually get synonymised into a single species name—becomes known to science. Species described earlier have had more time for the accumulation of synonyms. However, the difference between lag values and years remaining till present day indicates that for most of taxonomic history, this diversity characterising a species becomes known to science well before time becomes limiting (Figure 3a). Exceptions around present day violate the Principle of Priority and represent a very small fraction of the total species described around present day (Figure 2c). That species described earlier are wide-ranging (Collen et al., 2004; Gaston et al., 1995; Guedes et al., 2025; Higgs and Attrill, 2015), are more morphologically variable (Ficetola et al., 2016; Sides et al., 2014), and have more synonyms (Figure 3b) suggests a gradually changing trend in taxonomy of describing species encompassing smaller amounts of biological variability. In agreement, most species described in the latter half of the 20th century lack any synonym (Figure 2c).

Even though there are no cross-taxon trends of when the species accepted today were described (Appeltans et al., 2012; Figure 1), the proportion of accepted-names published each year remains initially constant across taxa (Figure 2a–b, Supplementary figure S1), suggesting that for this period in taxonomic history, the falsification ratio of a described species-name remains similar across taxa. While there is no discernable trend for synonymization for some fossil taxa like dinosaurs (Benton, 2008), our results show that given large sample sizes, there exists a simple and strong relationship for the temporal trends of synonymization across several extant taxa (Figure 3a–b).

The number of species with a lag time of zero peaks around 1900 (Figure 2d). These represent synonyms with a similar specific epithet as the accepted species-name. One major manner by which such synonyms with similar specific epithets arise is by new name combinations (Article 34, ICZN; “The Code Online | International Commission on Zoological Nomenclature”). That is, after the description of widespread, morphologically highly variable species, species described during this time were falsely allied to existing species or were falsely placed into distinct higher-level taxa (like the genus), with later rectification by taxonomic reassignment at higher levels of classification like the genus. For example, *Lacuna uchidai* (Habe, 1953) is a currently accepted mollusc species-name which was originally described as *Stenotis uchidai* (Habe, 1953) and arose as a new combination. This period is therefore characterised by species described for which accurate higher-level taxonomic relationships were not accurately determined during the time of description.

Next, because many recently-described species continue to be morphologically defined (Agapow et al., 2004) and formally described as allied to pre-existing species (Nerlekar et al., 2022), and have no synonyms (Figure 2c), they might represent a decrease in the amount of variability circumscribed within a species.Our results therefore depict gradually changing taxonomic praxis along with species concepts becoming narrower with time. While there are few studies directly quantifying the quality of species descriptions (Poulin and Presswell, 2016), our results instead suggest them via the novel analyses of trends of synonym-accumulation.

Notably, when a single species is split into multiple species upon taxonomic revision, synonyms of the former parent species may not be transferred to the newly erected species. For example, former deep-water populations of *Ophiacantha bidentata* Bruzelius, 1805 were split off into *Ophiacantha fraterna* Verrill, 1885 by most taxonomists based on morphological evidence. However, existing heterotypic synonyms of *O. bidentata* (*O. spinulosa* Müller & Troschel, 1842 and *O. arctica* Müller & Troschel, 1842) are not synonyms of *O. fraterna* (Martynov and and Litvinova, 2008). However, in spite of this non-transference of parental synonyms to newly split-off taxa, description date correlates with synonym number across taxa, datasets, timespans, and methods of quantification (Appeltans et al., 2012; Eggleton, 1999; Guedes et al., 2025), suggesting that the loss of synonym names in the aforementioned manner is unlikely to majorly impact our results.

Lastly, we note that while trends of synonymization have been used to estimate how many currently-accepted species shall later be falsified (Alroy, 2002; Costello and Wilson, 2011; Wortley and Scotland, 2004), our results instead depict the species concept itself changing with time.

## Conclusion

We use a comprehensive dataset of marine species and their synonyms and show that across taxa, while species description lacks discernible trends, the trends of synonymization are remarkably similar across taxa. These similarities suggest overall changes in the manner taxonomy was practised through time, with a reduction in the amount of variability characterising species on moving towards the present day.

## Acknowledgements

We gratefully acknowledge the comments and feedback of Akshay Bharadwaj, Arun Ravi, Balaji Chattopadhyay, and Dhrubojyoti Patra. We also acknowledge Devapriya Chattopadhyay for introductions from which this study emerged.

## Data and code availability

All data used are available from WoRMS (https://doi.org/10.14284/170). All codes used are available at the Zenodo URL https://doi.org/10.5281/zenodo.15285361.

## Author Contributions

VI: Conceptualisation (lead), dataset curation (supporting), formal analysis (equal), visualisation (equal), writing – original draft preparation (lead), writing – review and editing (equal).

AP: Conceptualisation (supporting), dataset curation (lead), formal analysis (equal), visualisation (equal), writing – original draft preparation (supporting), writing – review and editing (equal).

