## Supplementary Figures S1-S3 for "Species synonyms depict changing but taxon-independent taxonomic praxis"

Supplementary information

### Supplementary figure S1:

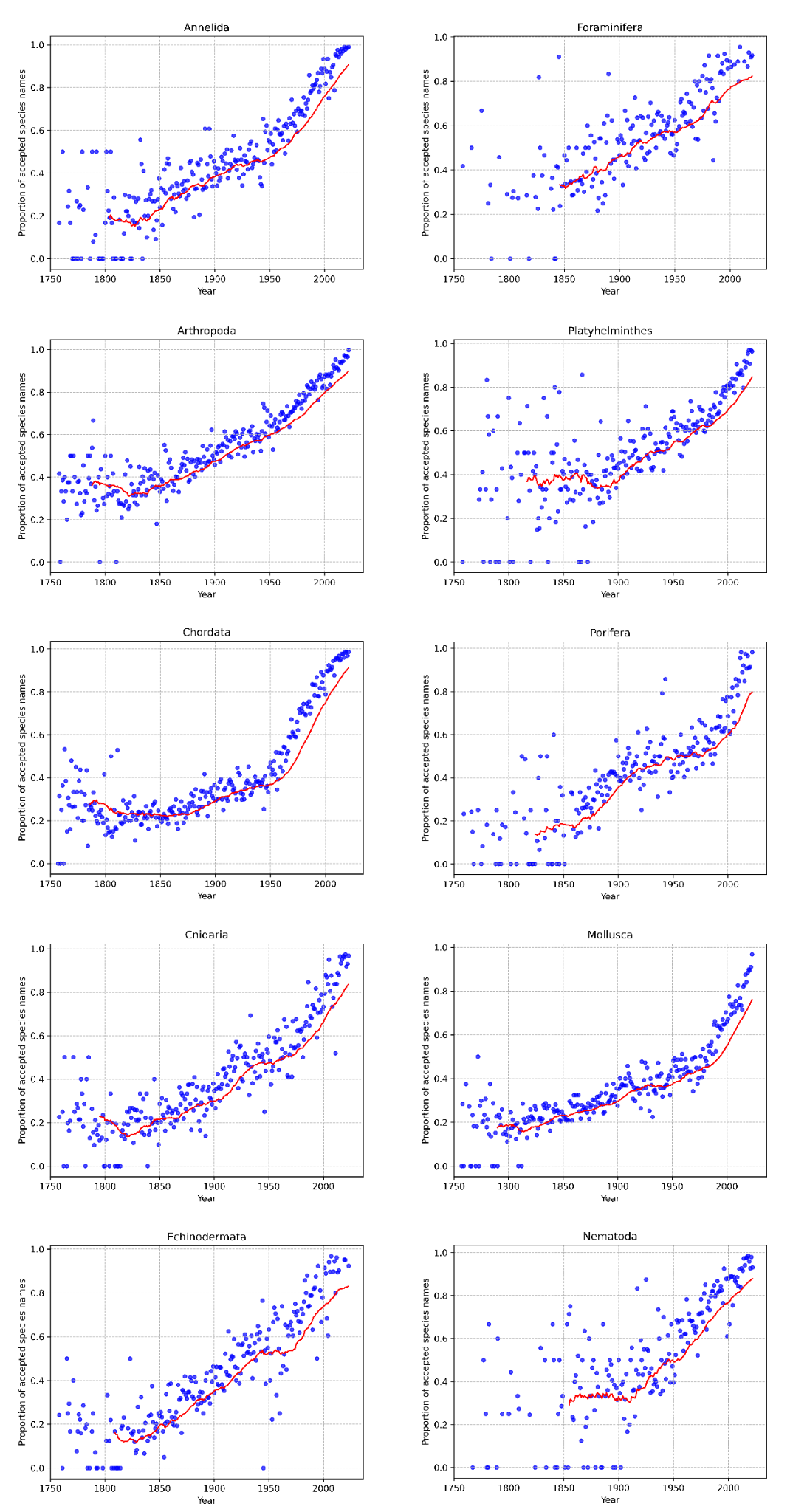

Figure S1:

Scatterplot depicting the fraction of all species-names published per year which are accepted today for all large phyla. Points represent values for a year and the red lines display the 20 year moving average.

Supplementary figure S2:

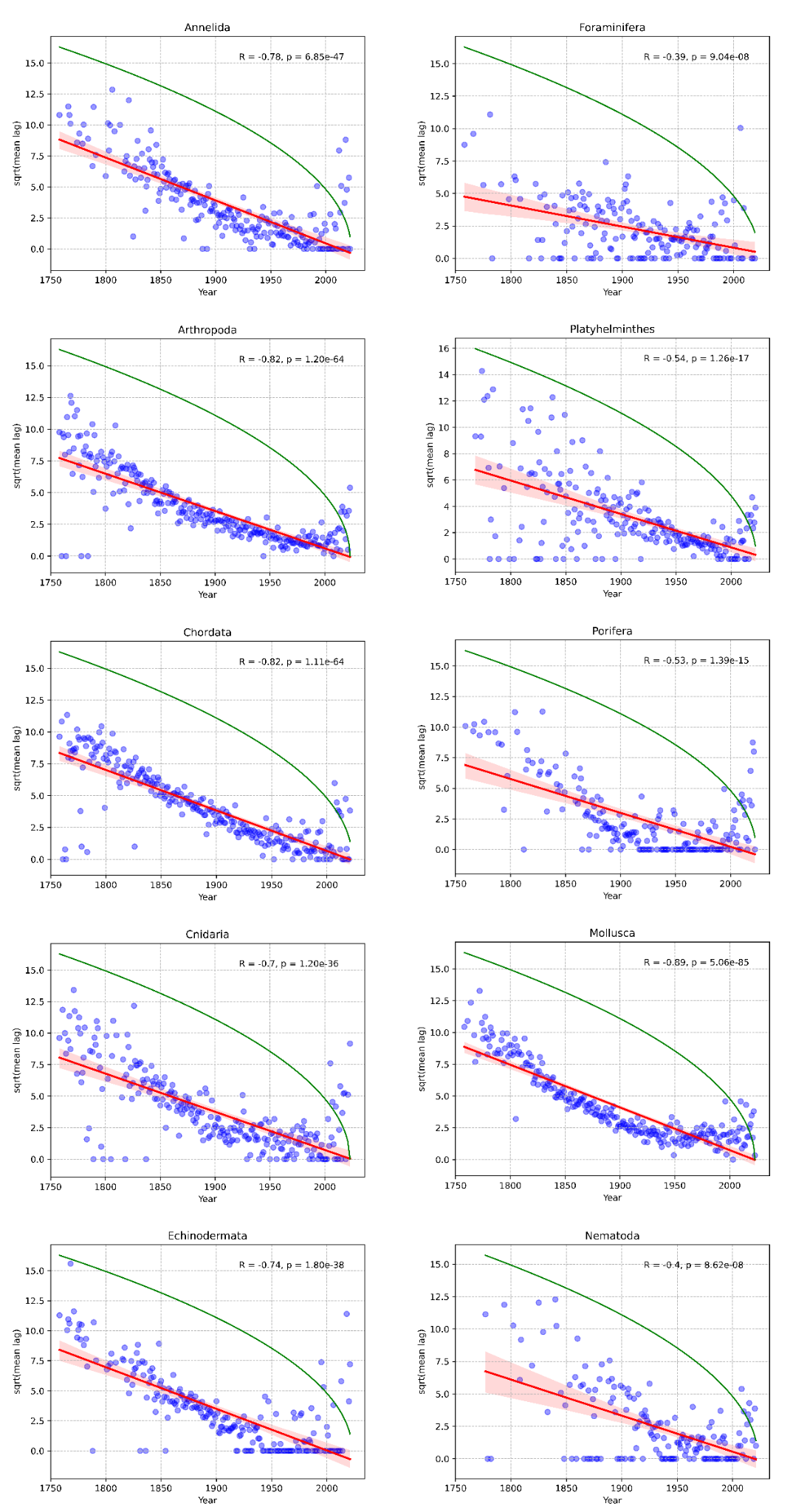

Figure S2:

Scatterplots depicting the mean lag value between the years of description of the newest synonym of the accepted species names published in a year for all large phyla. Points represent lag values. The red lines represent a straight line fit to the data and red bands indicate 95% confidence intervals. R = Spearman correlation coefficient. Lag values are square root transformed before plotting. The green lines represent the number of years till present day.

Supplementary figure S3:

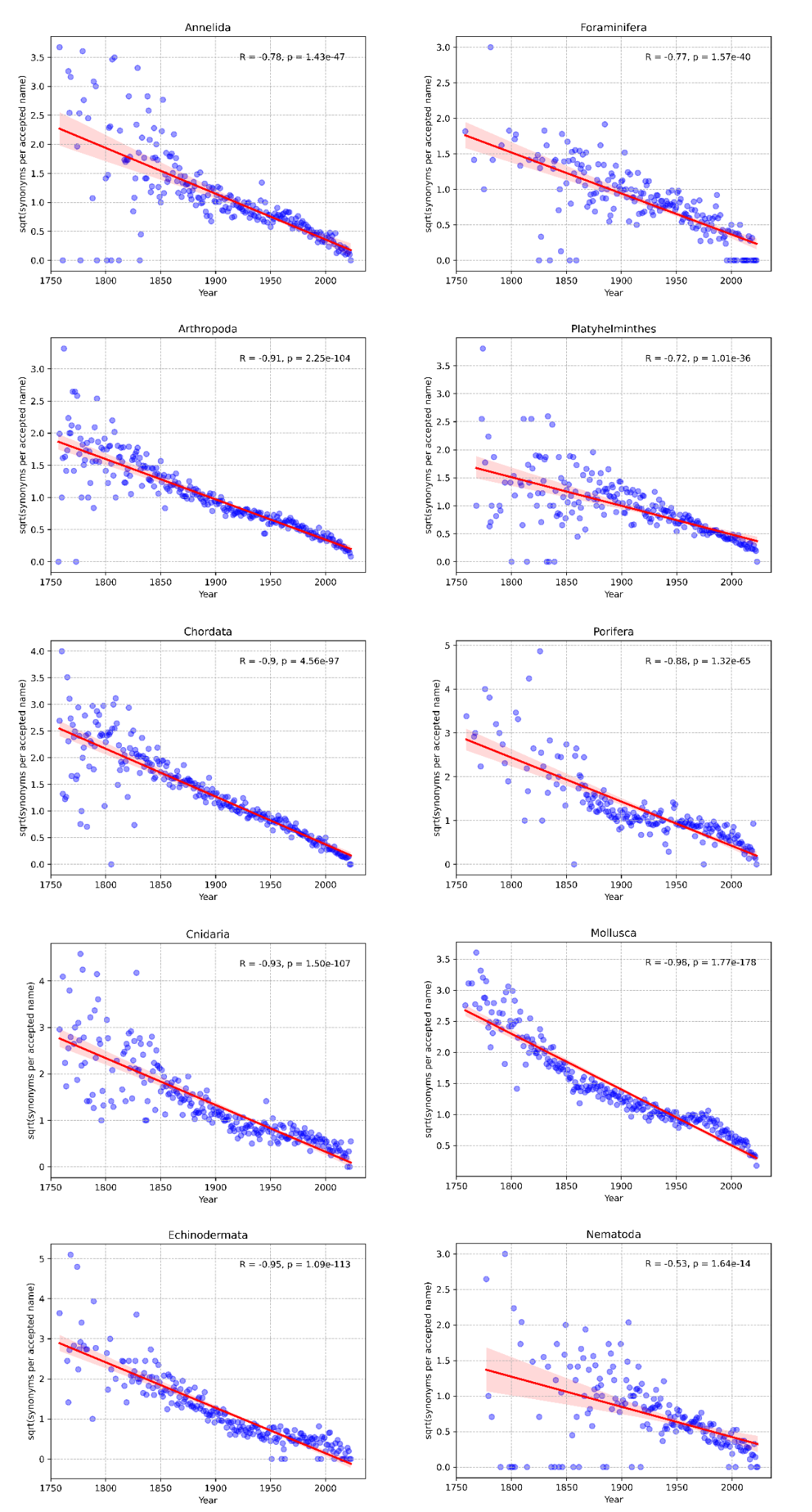

Figure S3:

Scatterplots depicting the mean number of synonyms for each accepted species-name published with time for all large phyla. Values are square root transformed before plotting. Points represent the mean number of synonyms. The red lines represent a straight line fit to the data and red bands indicate 95% confidence intervals. R = Spearman correlation coefficient.
